# What it takes to be at the top: The interrelationship between chronic social stress and social dominance

**DOI:** 10.1101/2020.06.29.177410

**Authors:** Merima Šabanović, He Liu, Vongai Mlambo, Hala Aqel, Dipesh Chaudhury

## Abstract

Dominance hierarchies of social animal groups are influenced by complex factors such as stress. Stress experienced by an animal prior to social interactions with a conspecific may be a determinant of their future social dynamics. Additionally, long-term occupancy of a specific hierarchical rank can have psychophysiological effects, leading to vulnerability to future stress.

The current study aimed to delineate differential effects of stress acting before or after hierarchy formation. Using the chronic social defeat stress (CSDS) paradigm we performed behavioural investigations to determine whether exposure to CSDS before hierarchy formation predicted the new dominance status. Moreover, in another study we investigated whether social rank predicted stress vulnerability.

We found that CSDS did not impede the establishment of dominance in new hierarchies as both stress-susceptible (socially avoidant) and –resilient (social) mice were able to attain dominant ranks. In contrast, within newly established hierarchies of stress-naïve mice, the subordinate, but not dominant, mice exhibit significantly greater avoidance of novel social targets. However, following exposure to CSDS, both lowest- and highest-ranked mice exhibit strong susceptibility to stress as measured by decreased interactions with a novel social target.

These results suggest that the response to chronic social stress did not determine social rank in new cohorts, but low-status mice in newly established groups exhibited lower sociability to novel social targets. Interestingly, exposure of a hierarchical social group to chronic social stress led to stress-susceptibility in both high- and low-status mice as measured by social interaction.

**Highlights:**

- Stress susceptibility to chronic social defeat did not impede the establishment of dominance in new hierarchies.
- Subordinate mice exhibit reduced social preference after hierarchy formation.
- Following chronic social defeat stress, both subordinate and dominant mice exhibit susceptible-like reduction in social interaction, but dominant mice exhibit the greater decrease in social preference as compared to baseline.

## 1. Introduction

Formation of dominance hierarchies is recognized as a universal and fundamental organizing mechanism for social animal groups [1]. Where resources are limited, social hierarchies determine an individual’s access to food, territory, or mating partners, and are readily formed due to their adaptive power of minimizing fighting among conspecifics living in close proximity [2]. Hierarchical rank has extensive effects on physical and mental health [3–5] and therefore could become maladaptive due to the risk factors associated with living in a particular rank. Previous research has shown extensive effects of rank on behaviour, including reproductive success [6], anxiety [7] (but see [8]), social motivation [9] and social contact [10], as well as on gene expression [7] and receptor expression [11]. During the formation of social hierarchies, ranks are not determined solely by intrinsic attributes, such as body size and weight, but are also affected by the environment and prior experiences of the animals. Chronic pain [12], stress [13–15] and sleep [16] can have marked effects on social behaviours and dominance hierarchies. Evidence suggests that stress has a complex link to social hierarchies as it could contribute to hierarchy formation as well as arise because of hierarchy maintenance. Laboratory rodents are a particularly pertinent model organism for investigating hierarchy formation as they are very social animals and allow for studies of neuronal mechanisms underlying behaviour. Both in the wild and in the lab, dominance hierarchies are readily observable due to the distinct patterns of behavioural characteristics in the different social ranks (reviewed in [17]). One of standard tests of dominance in mice is the competitive exclusion task or the “tube test” [18], first developed to study dominance differences between inbred strains [19]. This dyadic test offers clear and binary scoring of dominance, based on the use of space resources, that would otherwise be difficult to assess directly in the home cage.

Extensive effects of stress are found on a cellular, behavioural and physiological level as evidenced by alterations in synaptic plasticity [20], learning and memory (reviewed in [21]), hormonal responses [22] and sleep [23]. Nevertheless, the effects and causes of stress in relation to social hierarchies are poorly understood. For example, it is known that prior acute stress exposure renders an animal more likely to be in a long-term subordinate status after a conflict encounter [24]. Similarly, chronic stress such as restraint stress was linked to decreased display of social dominance in the tube test [13]. Thus, considering the social defeat stress (CSDS) paradigms based on the resident-intruder aggression which increases defensive and submissive behaviours in the test (intruder) animals [25], we hypothesized that animals susceptible to CSDS would be more prone to subordinance in subsequent hierarchy formation. In addition to the effects of prior stress exposure on hierarchy formation social rank may also affect susceptibility to future stressors. For example, a recent paper had shown that dominant mice were more susceptible to developing depression-like behaviours following CSDS [26, see 27 for review]. However, depressive-like phenotype can also be induced by long-term subordination alone [28]. Thus, we aimed to elucidate social hierarchy formation in male C57BL/6 mouse before and after exposure to stress.

We first investigated whether chronic stress differentially affected hierarchy formation, anxiety levels and diurnal locomotor rhythms in mice resilient or susceptible to CSDS. Our findings that resilient or susceptible mice equally exhibit dominant or subordinate status suggests that processes that lead to stress-resilience or -susceptibility do not affect hierarchy formation. Moreover, anxiety and diurnal locomotor rhythms were not affected either after stress or hierarchy formation. We next investigated whether establishing and maintaining a particular social rank, prior to any further stress exposure, could be stress-inducing. Prior to stress, baseline sociability was shown to be an indicator of social rank since subordinate mice exhibited significantly less social interaction compared to the dominant mice to a novel social target. We also observe that dominant mice exhibit the greatest decrease in social preference following CSDS. However, in contrast to previous reports [26] dominant mice were not found to be more susceptible to CSDS when measured on a social preference test since the distribution of social preference scores post CSDS was similar in dominant and subordinate mice. Moreover, no rank-dependent differences in anxiety, anhedonia, or wheel-running activity, were found. Our observations suggest that social rank status may not be predicted by resilience or susceptibility to stress as measured by social interaction.

## 2. Methods

### 2.1 Ethics

All animals were kept in Institutional Biosafety Committee (IBC) approved housing at the New York University Abu Dhabi (NYUAD) animal facility. All experimenters completed the Collaborative Training Initiative (CITI) Animal Care and Use Course, which meets United States Department of Agriculture (USDA) and Office of Laboratory Animal Welfare (OLAW) criteria for training in the humane care and use of animals in research. All animal protocols were in accordance with the National Institute of Health Guide for Care and Use of Laboratory Animals (IACUC Protocol: 150005A2) and have been approved by the NYUAD Animal Care and Use Committee.

### 2.2 Animal rearing

All experiments were performed on male C57BL/6J mice (Jackson Laboratory). Behavioural experiments commenced when animals were 7 weeks old and were completed by 15 weeks of age. Retired male CD1 breeders (Charles River Laboratory) were used as resident aggressors during the CSDS paradigm. All mice were maintained under standard housing conditions at a humidity of 50±10%, temperature of 23±2°C and a 12h light/dark cycle (7AM-7PM), with *ad libitum* access to food and water. Wood shavings were used as enrichment in the home cage with social housing, unless isolation was required by the experimental protocol.

### 2.3 General experimental design

Upon arrival, mice were ear-marked and allowed one week of acclimation before onset of experiments. Animals were weighed weekly to ensure healthy weight and ensure weight-matching within cage groups. All behavioural tests were conducted during the light period of higher activity (2PM-7PM), and the mice were habituated to the recording room and lighting conditions for at least 30min prior to testing. Between animals, the behavioural apparatus was cleaned with MB-10 solution (active ingredients: 20.8% sodium chlorite and 7.0% sodium dichloroisocyanurate dehydrate) for disinfection and elimination of olfactory cues. Mice tested for the effect of stress on subsequent hierarchy formation were single housed upon arrival to avoid confounding effects of prior hierarchical rank. Within the experiment, variance was reduced by using all male mice that were age- and weight-matched per cage. The same female experimenter handled mice prior to and during testing whenever possible to reduce stress and anxiety responses.

Please take note that all mice used in the study of the effects of rank on stress-susceptibility underwent virus injection surgeries for projection tracing (data not shown). These surgeries were performed at 6-week-old mice who were allowed one week of recovery before re-housing into novel weight-matched groups of four. All animals recovered well from anaesthesia and the surgery, were healthy and displaying normal behaviour. We therefore believe the surgery did not confound the experimental results reported here.

### 2.4 Social dominance tests

A minimum of two weeks of cohabitation were allowed for stable hierarchy formation, as defined by earlier studies [8]. The validity of the tube test was critiqued based on whether it is a true measure of dominance considering the possible confounding effects of sensorimotor capacity, learning ability and spatial context [29,30]. Therefore. we have used a battery of dominance tests displaying high consistency of ranking results with those obtained by the tube test, validating that dominance is indeed the underlying variable being measured in the context of our study. The rank was established first in the tube test, and then followed by three supplementary dominance tests. We have only used male animals in this study as female mice are not commonly used in social dominance assessments based on territoriality and vocalizations since they rely more on intrinsic attributes and social feedback to establish a hierarchy rather than prior social experience [31]. However, some studies were able to show stable linear hierarchies in female mice but the effect of oestrus stage has to be taken into account [32].

#### 2.4.1 Social confrontation tube test (TT)

Our customized automated TT system (Clever Sys Inc.) consists of a clear Plexiglass tube 55cm in length and 2.5cm in diameter, sufficiently wide for one mouse to walk through but not for two mice to pass each other. The tube is connected to a 10 × 10cm box on each side. The box in which a mouse is initially placed is called the “starting box”, for which the box at the other end would be the “goal box”. Automated doors are placed at the box exits and in the middle of the tube. During the two-day training phase, each mouse was given 10 trials, 5 starting at each side, in order to learn to enter the tube, initiate the middle door to open when within 3cm of reaching it, and pass to the goal box. If the mouse remained stationary for longer than 5s, or began retreating, a gentle push from behind was used to direct movement towards the middle door. The use of food deprivation and food reward previously showed no effect on animals’ motivation to complete the task [18] and was therefore not used in this study.

Our pilot experiments used a 7-day testing phase, but the animals started displaying signs of stress and reduced task motivation in the later days, so we opted for 4-day testing instead. On each day of the testing phase, each mouse explored the tube once from each side prior to starting the confrontation trials. Using a round-robin design, all pairs of mice from the same cage were tested (6 pairs per social group of 4 mice). During the trial, both mice were guided into the tube simultaneously from their respective starting boxes. The starting side for each mouse alternated between trials. When both mice were within 3cm of the middle door, the door opened, and the social confrontation trial began. The trial ended when one of the mice retreated with all four paws to its starting box, therefore becoming the “loser” or the subordinate (**Fig.1A**). The mouse that forced its cage mate to retreat was termed the “winner” or the dominant. In between trials, mice were kept in separate clean holding cages. The confrontation trials were repeated for four consecutive days with the randomized order of the pairs and the cages. The experimenter remained stationary during each trial in a designated position in the room to maintain cue consistency.

**Fig. 1.**
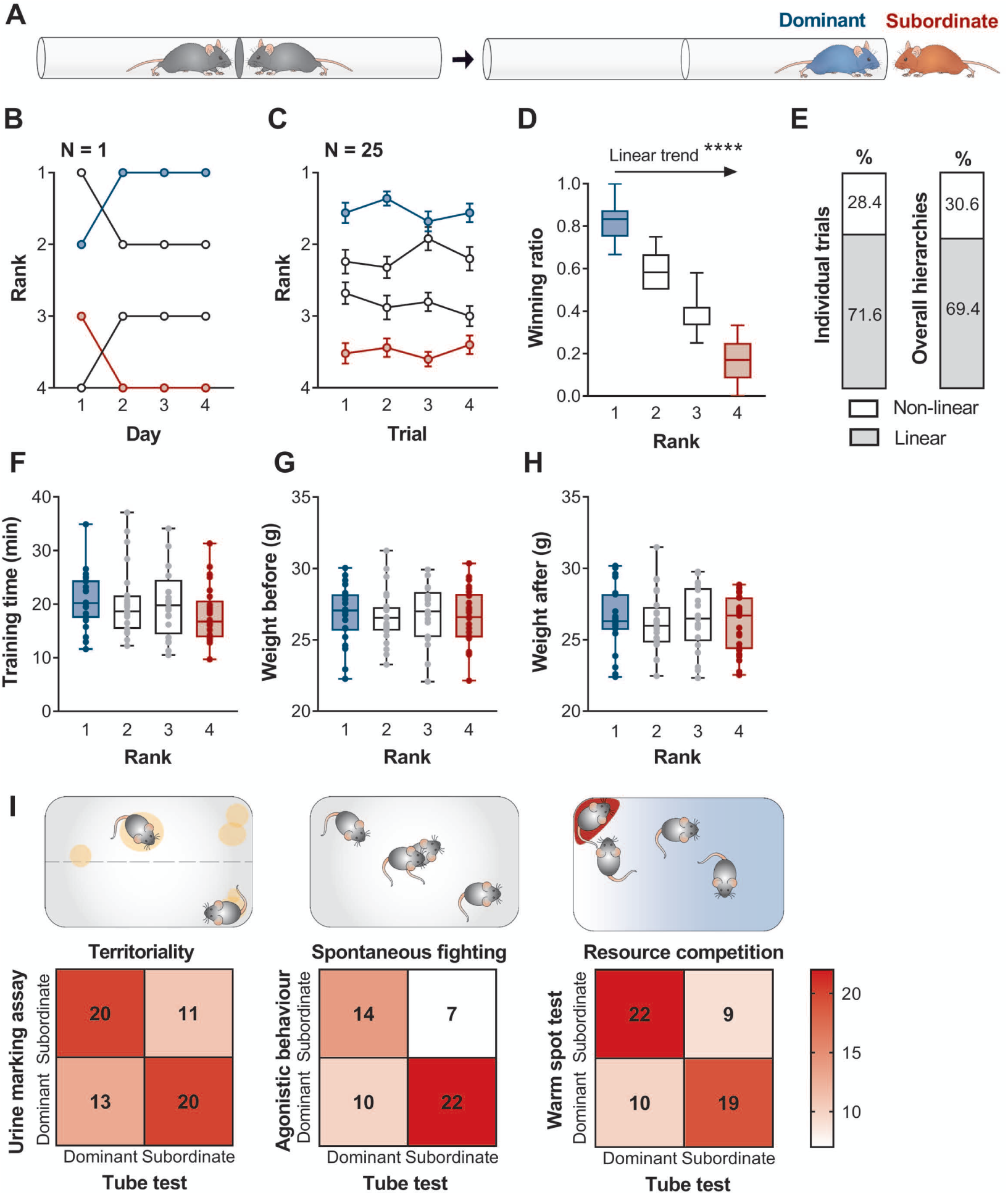
Social confrontation tube test was validated to yield predominantly linear and stable hierarchies consistent with other measures of dominance. (A) Diagram of a confrontation trial. The subordinate mouse retreats from the tube first. (B) Representative image of the rank stability in a cage of four mice over the four days of testing. (C) Overall rank stability shows the average rank of animals belonging to each rank group (as determined at the end of TT) calculated for each day of testing. (D) Winning ratio of ranked mice (N=32-36 per rank) shows a linear trend from the most dominant to the most subordinate. (E) Individual daily trials and final hierarchies are predominantly linear. (F) The total time spent in the tube during a two-day training phase does not differ between ranks (N=20-25 per rank). (G) Weight is not a factor in establishing dominance as groups are weight-matched prior to TT (N=21-25 per rank). (H) Weights of ranked mice are comparable after testing (N=21-25 per rank). (I) TT correlates with rankings from three other dominance tests: urine marking assay, UMA (p=0.0509); agonistic behaviour test, ABT (p=0.0229); and the warm spot test, WST (p=0.0090). Data are shown as mean ± SEM or box plot with whiskers denoting the min. and max. values. ****p<0.0001.

The winning ratio was calculated as the number of all trials won by that mouse divided by the total number of trials. This determined the index of overall dominance where rank 1 and rank 2 mice (winning ratio > 0.5) were the dominant, while rank 3 and rank 4 mice (winning ratio < 0.5) were the subordinate mice.

#### 2.4.2 Territory urine marking assay (UMA)

Mice are territorial animals and urinary scent marking serves to indicate territorial boundaries and dominance status, strongly influencing their aggressive interactions [33]. The number of scent marks can predict both aggression scores and social dominance status in mice [34]. Using the round-robin design, each of the six possible pairs from the same home cage was tested (two pairs per day for the total of three days) following a protocol established by Wang et al. [35]. The number, size and the distance of urine marks from the central partition were scored blinded to the tube test result. “Dominant” males were identified as those making more urine marks and/or close to the partition, whereas “subordinate males” were those urinating in fewer locations and/or further away from the partition. A total of 20 cages of animals were tested, with 16 yielding unambiguous dominant-subordinate relationships within the group.

#### 2.4.3 Warm spot test (WST)

This test was adapted from Zhou et al. [36]. A rectangular plastic cage 29.5 × 18cm was placed on ice, cooling the floor of the cage to 0-4°C. The mice were first habituated to the cold cage for 30min. Then, they were transferred to a new cold cage with a 5 × 5cm warm pad heated to 34°C. Correct temperatures were ensured by monitoring with an infrared thermometer. As the warm spot was big enough to permit the stay of only one adult mouse, the competition of the four mice for the warm spot was videotaped for 20min and the time each mouse spent occupying the warm spot was analysed. Dominance, characterized by longer warm spot occupation times, was scored blinded to the tube test result.

#### 2.4.4 Agonistic behaviour test (ABT)

Mice group-housed together for an extended period will not exhibit extensive aggressive behaviour toward each other. Others have reported that agonistic behaviour is potentiated upon placing the animals in a new cage which requires the animals to claim the new territory [35]. We observed increased instances of agonistic interactions immediately after returning the animals to their home cage after behavioural testing, presumably due to the need to reinforce their status upon re-entering their territory. Accordingly, tail-marked mice were videotaped for 15min upon returning to the home cage following either the UMA or the WST, recording the occurrence of spontaneous fighting and offensive or defensive behaviour. Offensive behaviours were characterized as chasing and attacking, while the submissive behaviours included flight, freezing and submissive posture (exposed abdomen, limp forepaws and head angled up). In most cases, only one mouse out of four in the group would initiate an attack. Agonistic behaviour was observed in 13 out of 23 cages tested.

### 2.5 Inducing and characterizing chronic social stress

#### 2.5.1 Chronic social defeat stress (CSDS)

The paradigm was adapted from Golden et al. [37] to a duration of 15 days. CD1 mice were screened for aggression in their home cage and rescreened prior to starting CSDS, excluding non-aggressive mice. An experimental mouse was placed into the home cage of a CD1 aggressor mouse for 10min during which time it endured several bouts of physical attacks by the aggressor. The CD1 and experimental mouse were then maintained in sensory contact for 24h using a perforated plexiglass partition dividing the resident home cage in two. On each consecutive day, the experimental mice were exposed to a new CD1 mouse home cage to avoid habituation to the aggressor. The repeated social defeats were performed between 3-5PM. Control mice were housed in pairs within a cage setup identical to that of CSD mice, with two mice continuously separated by the perforated Plexiglass divider. Control animals were removed from the room during the defeat sessions, to avoid exposure to stress-induced vocalizations by their conspecifics. 24h after the last defeat session, the mice were taken out of the CSD cages and single housed in new cages, allowing a minimum of 3h of habituation before starting the social interaction test.

#### 2.5.2 Social interaction test (SI)

A two-stage SI test was adapted from Golden et al. [37]. In the first 2.5min-long non-social session (no social target present), the mouse could freely explore a square-shaped arena (42 × 42cm) containing a clear Plexiglass cage with a wire mesh (10 × 6.5cm) placed on one side of the arena. In the second 2.5min-long social session (with a neutral novel social target present), the experimental mouse was reintroduced back into the arena with an unfamiliar CD1 mouse contained behind a wire mesh cage. Between the non-social and social sessions, the mouse was removed from the arena and placed in his home or neutral cage. Video tracking software (TopScan, Clever Sys Inc.) was used to measure the amount of time the experimental mouse spent in the “interaction zone” (24 × 15cm surrounding the wire mesh cage), “corner zone” (9 × 9cm from opposite walls), as well as the total distance travelled by the mouse. The social interaction ratio (SI ratio) was obtained by dividing the time spent in the interaction zone in the social session divided by the object session. Susceptibility to stress was characterized by a reduction in SI ratio to values below 1.0, indicating social avoidance. Two separate populations are defined as “stress-susceptible” (post-CSDS SI ratio <1.0) and “stress-resilient” (post-CSDS SI ratio >1.0). Social interaction deficits are transferrable across species, observed with an unfamiliar CD1 as well as C57 social target [14].

#### 2.5.3 Sucrose preference test (SPT)

Animals were single housed and habituated to two bottles of 1% sucrose for two days, followed by a 24h-period of food and water deprivation. In the 3h test period, the animals were given one bottle of 1% sucrose and one bottle of water, with bottle positions switched halfway through the experiment, to control for any side-preference. The sucrose and water bottles were weighed before and after the test, recording the total consumption of each liquid. Sucrose preference was defined as: total sucrose consumption / total liquid consumption (water and sucrose).

#### 2.5.4 Wheel-running assay

Voluntary wheel-running cages were placed in circadian cabinets (Phenome Technologies) with a maximum of 6 cages per row. The light and temperature of the chambers was controlled by the ClockLab Chamber Control software (ACT-500). The running wheel cages are available from Actimetrics (model: ACT-551-MS-SS) and consist of a Tecniplast model 1144B cage bottom (33.2 ⨯ 15 ⨯ 13cm) and a wire bar lid. The wheel is stainless steel, 11cm inside diameter, 5.4cm wide, with 1.2mm wide bars placed 7.5mm apart. The infrared (clickless) sensor clips onto the lip and rail of the cage and detects the spokes of the wheel passing by. The sensor was connected via cable to the ClockLab digital interface (ACT-556). ClockLab Data Collection software (ACT-500) registered each revolution of the wheel as a count. The number of counts per minute of each wheel was recorded and the final analysis was done on the total counts per hour with ClockLab Analysis Version 6 (ACT-500). The total of one week of recording was used.

The running wheels are commonly used as a measure of circadian activity rhythms, but evidence suggests that wheel-running is also rewarding to rodents (see [38] for review) and can therefore potentially be used as a surrogate for motivated physical activity. Some studies suggest voluntary wheel-running has anti-depressive and anti-anxiety-like effects, but the review of previous research shows that a minimum of 3-4 weeks of unrestricted access are required for such effects to be significant, [38]. Considering this protocol includes only one week of wheel-running, we do not expect that such behavioural alterations presented a significant confounding variable for the experiments that followed the wheel-running activity assay.

### 2.6 Anxiety tests

All animals were handled for a minimum of two days prior to onset of baseline anxiety measurements, allowing the mice to habituate to the experimenter interaction. TopScan (Clever Sys Inc.) video tracking system recorded the time spent in each zone, as well as bouts of entering each zone and total distance travelled as measures of exploration.

#### 2.6.1 Open field test (OF)

The apparatus consists of the same arena used for the SI test (42 × 42cm), with the “center” zone defined as the inner 32 ⨯ 32cm. Under red light, mice were placed into the center of the arena and allowed 15min of free exploration. Thigmotaxis in the OF was defined as the percentage of testing time the animal spent near the walls of the arena and not in the center.

#### 2.6.2 Elevated plus maze test (EPM)

A grey polyvinyl chloride (PVC) apparatus was in the “+” configuration comprising of two open arms (34 ⨯ 6cm) perpendicular to two closed arms (34 ⨯ 6cm with 21.5cm tall walls) with a center zone (6 ⨯ 6cm). The entire apparatus was 60cm above the ground and illuminated by red light. The animal was placed in the center zone, opposing the experimenter, and allowed 5min of free exploration. Thigmotaxis in the EPM was defined as the percentage of testing time the animal spent in the closed arms of the maze.

### 2.7 Statistical analyses

Animals were randomly assigned to treatment groups, but cage groups were matched by weight. Where possible, the experimenter was blinded to treatments. Statistical analyses were performed using GraphPad Prism software v7. All values are given as a mean ± SEM. All statistical tests were two-tailed and the significance was assigned at p<0.05. The D’Agostino-Pearson omnibus normality test and Brown– Forsythe test were used to test normality and equal variances between group samples, respectively. When normality and equal variance between sample groups was achieved, ordinary one-way ANOVA (followed by Tukey’s multiple comparisons test), repeated measures two-way ANOVA, unpaired or one sample t-tests were used. Where normality or equal variance of samples failed, Mann–Whitney U test, Kruskal-Wallis test with Dunn’s post-hoc multiple comparison, or Wilcoxon signed rank test was performed. Linear regression and Fisher’s exact tests were used for correlation and contingency analyses.

## 3. Results

### 3.1 Hierarchy formation and TT validation

Following two weeks of agonistic activity, establishment of social hierarchies among cagemates was determined in a battery of social dominance tests. In the TT, the measured hierarchies were consistent over the four-day testing period, both within and across groups respectively (**Fig.1B-C**), and exhibited a linear trend from the most dominant to the most subordinate animal (**Fig.1D**: One-way ANOVA F(3,132)=378.4, p<0.0001: post-test for linear trend slope= -0.2171± -0.0064, R^2^=0.8956, F(1,132)=1135, p<0.0001). In a linear hierarchy model, the top individual (“alpha”) dominates over all others. Each subsequent rank is singly occupied, down to the most subordinate mouse (“omega”) that is dominated by all other members. Nearly 70% of hierarchies observed in this experiment were linear (**Fig.1E**), but other hierarchical structures were observed, such as non-transitive and despotic (one alpha with other members not having clear ranks). The TT rank is not induced by the testing procedure, as the time spent in the tube during the training phase was not an indicator of success during testing (**Fig.1F**: Kruskal-Wallis H(3)=3.83, p=0.2804) and neither was the weight profile before (**Fig.1G**: One-way ANOVA F(3,88)=0.3467, p=0.9576) or after testing (**Fig.1H**: One-way ANOVA F(3,88)=0.1442, p=0.9331). The validity of TT-obtained ranks is supported by correlation with other dominance measures that highlight different manifestations of dominance behaviour. Dominance ranks from the TT were consistent with ranks obtained via three other methods: territoriality in the UMA, spontaneous fighting in the ABT, and resource competition in the WST (**Fig.1I**: Fisher’s exact t-tests (2-sided) UMA p=0.0509, ABT p=0.0229, WST p=0.0090). The degree of correlation ensures that dominance is the common underlying factor being measured. Ranked mice belonged to two experimental groups based on exposure to CSDS before or after hierarchy formation, as described in sections 3.2 and 3.3.

### 3.2 Experiment 1: Effect of CSD on novel hierarchy formation

After a baseline SI test, one cohort of animals underwent 15 days of CSDS, followed by another SI test, as well as SPT and 1-week wheel-running assay. The mice were then housed in weight-matched groups such that there is one CSDS-stressed mouse together with three stress-naïve controls in one cage (total of N=19). After two weeks of hierarchy formation, TT and supporting dominance tests were performed as described previously. Anxiety was measured prior to starting and after completing behavioural tests. The full timeline is shown in **Fig.2A**.

**Fig. 2.**
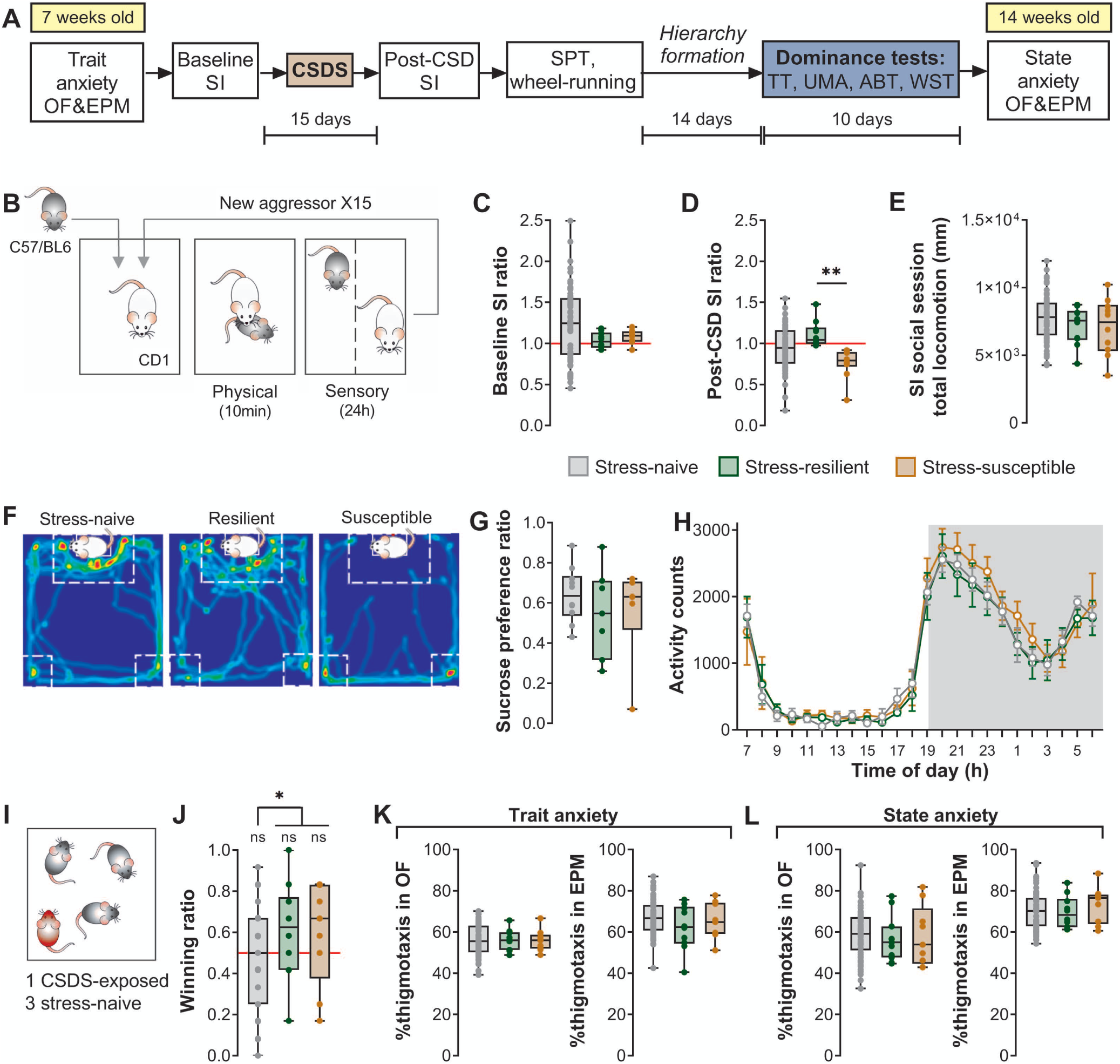
CSDS results in two distinct populations based on social interaction profiles that did not differ in subsequent hierarchy formation. (A) Timeline of behavioural studies investigating the effect of chronic stress on subsequent hierarchy formation. The age of mice is given in yellow boxes. (B) 15-day CSDS paradigm consisted of daily sessions of 10min physical stress followed by 24h of sensory stress. (C) Before chronic stress exposure, SI ratios did not differ between groups. (D) After CSDS, a subset of mice termed “stress-susceptible” exhibited an SI ratio lower than that of “stress-resilient” mice. (E) Level of exploration, as measured by total distance travelled during the social session of the SI test, was comparable across stress groups. (F) Representative trace of the time spent interacting with a social target shows that stress-susceptible mice avoid the interaction zone around the social target mouse and escape to the corner zones. (G) None of the groups exhibit anhedonia following CSDS. (H) All groups exhibit similar daily wheel-running activity profiles. (I) One CSDS-exposed mouse was group-housed with three stress-naïve controls. (J) There was no difference in dominance between stress-exposed groups, but CSDS-exposed mice display a more dominant status on average than their stress-naïve cagemates. (K-L) Anxiety profiles in both OF and EPM anxiety tests did not differ between stress groups either before or after behavioural testing. N_Stress-naïve_=60, N_Resilient_=N_Susceptible_=10. Data are shown as mean ± SEM. **p<0.01. CSDS: chronic social defeat stress. SI: social interaction. SPT: sucrose preference test TT: tube test. UMA: urine marking assay. ABT: agonistic behaviour test. WST: warm spot test. OF: open field. EPM: elevated plus maze.

#### 3.2.1 CSDS induces reduced social preference in a subset of stress-susceptible mice

Chronic social stress was induced using the CSDS procedure based on social conflict, as described in section 2.5 (**Fig.2B**). In the baseline SI test, mice exhibit a normal distribution for social preference (**Fig.2C**: One-way ANOVA F(2,77)=1.498, p=0.2300). Following CSDS, animals could be distinguished as belonging to two separate groups – those rendered stress-susceptible (socially avoidant), recognizable by their lower SI ratios (<1.0) that were significantly different from the SI ratios of stress-resilient mice (**Fig.2D**: Kruskal-Wallis H(2)=11.23, p=0.0036; Dunn’s multiple comparisons naïve-resilient p=0.1095, naïve-susceptible p=0.0655, resilient-susceptible p=0.0024). This reduced SI ratio is not due to reduced exploration (**Fig.2E**: One-way ANOVA F(2,77)=0.9374, p=0.3961) but rather reflects the tendency of stress-susceptible mice to avoid the interaction zones of the SI arena (**Fig.2F**). Reduced social preference in this case may not be a marker of a depressive-like phenotype as mice do not exhibit anhedonia in the SPT (**Fig.2G**: One-way ANOVA F(2,20)=0.5641, p=0.5776; Fig.A2D) or aberrant wheel-running activity (**Fig.2H**: Two-way RM ANOVA stress group effect F(2,20)=0.3953, p=0.6786). Accordingly, “stress-susceptibility” was therefore used as a measure of stress-induced social avoidance.

#### 3.2.2 CSDS did not diminish success of stress-exposed mice in subsequent hierarchy formation

Following CSDS, the mice were weight-matched and group-housed such that there is one stress-resilient (N=10) or stress-susceptible (N=9) mouse together with three stress-naïve controls (total N=57) per cage (**Fig.2I**). After two weeks of hierarchy formation, winning ratios obtained in the TT were compared between stress groups. Surprisingly, neither stress-resilient nor stress-susceptible mice were more likely to be subordinate and their average winning ratios were comparable (**Fig.2J**: stress-naive Wilcoxon Signed Rank median=0.5, p=0.2239; One sample t-test, resilient t(9)=1.423, p=0.1885 and susceptible t(8)=1.214, p=0.2594). Our post-hoc exploratory analyses suggest that CSDS-exposed mice occupy the more dominant positions in their respective cohorts (**Fig.2J**: stress-naïve vs. CSD-exposed Mann-Whitney U=365, exact p-value=0.0330), but more rigorous follow-up studies with higher power would be needed for conclusive results.

#### 3.2.3 Neither trait nor state anxiety could be used as a predictor of stress-susceptibility

OF and EPM anxiety tests were performed prior to starting behavioural manipulations to establish the characteristic of the individual (trait anxiety). Moreover, OF and EPM tests were performed after exposure to the CSDS and dominance test to determine the effects of experiencing chronic stress and hierarchy formation on anxiety (state anxiety). Both OF and EPM tests use the measure of thigmotaxis, defined as the tendency to remain close to walls or enclosed spaces, as a proxy for high anxiety. All experimental groups remained comparable to each other at both time-points (One way ANOVAs, **Fig.2K**: OF: F(2,73)=0.03089, p=0.9696; EPM: F(2,73)=1.332, p=0.2704; **Fig.2L**: OF: F(2,73)=0.2323, p=0.7933; EPM: F(2,73)=0.4725, p=0.6254). Explorative behaviours were not affected since behavioural measures such as bouts of zone entries and total locomotion were consistent between groups in all tests (**Fig.S1**).

### 3.3 Experiment 2: Effect of social status on stress susceptibility

To delineate whether newly formed hierarchies would predispose a certain rank to greater stress susceptibility, a second cohort of mice were first matched by weight and group-housed immediately after trait anxiety tests (total of N=16 groups). After ranks were determined in dominance tests and a baseline SI was recorded, all ranks underwent CSDS with a random small subset of mice serving as CSDS-controls (and therefore being excluded from further comparison based only on CSDS-exposed mice). The same tests of stress were performed as in Experiment 1, followed by state anxiety recordings. The experimental timeline is shown in **Fig.3A**. To minimize the variability due to different hierarchical structures, main comparison was limited to the clear alphas or omegas of a group, the rank 1 and rank 4 mice respectively.

**Fig. 3.**
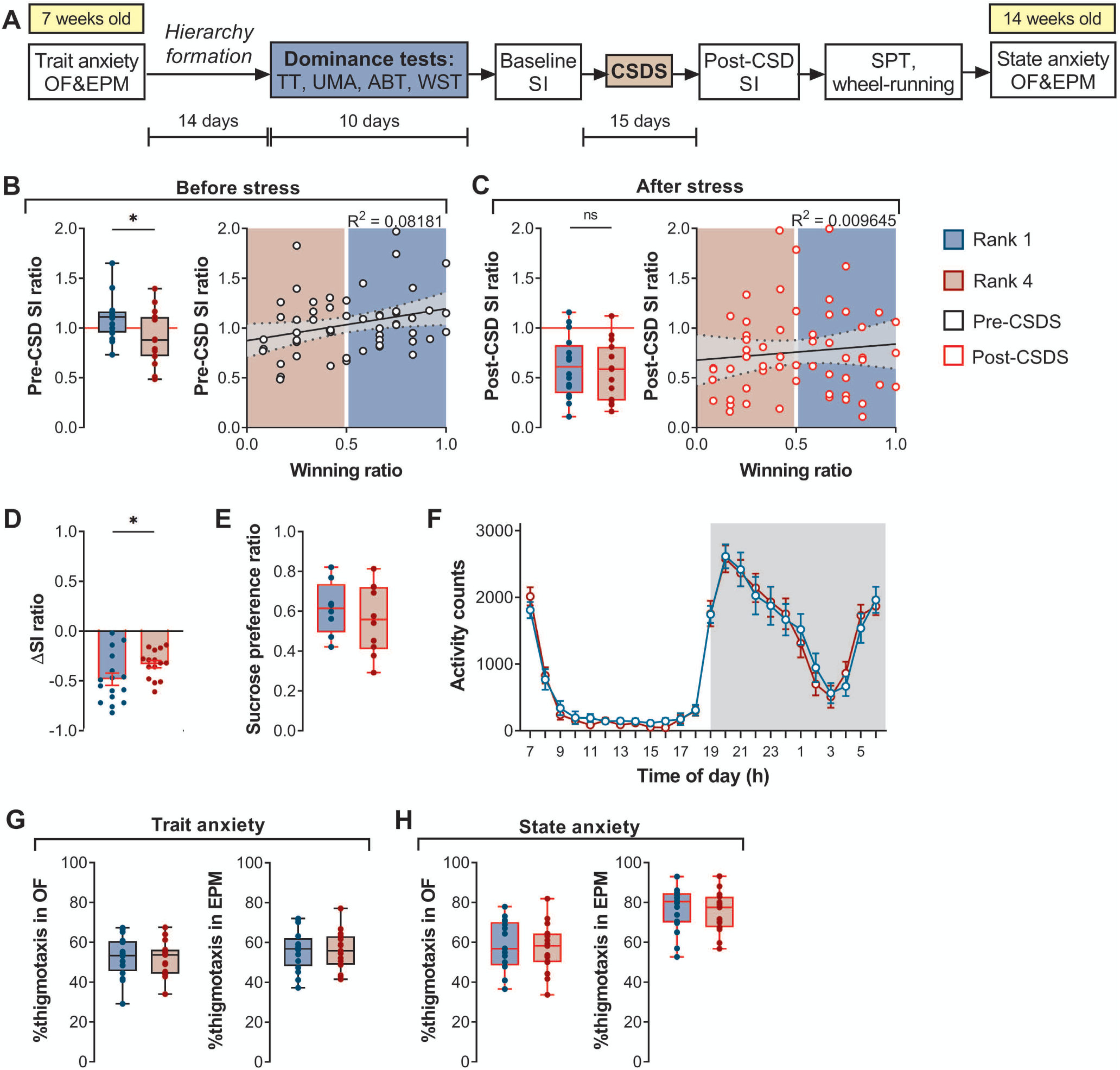
Baseline, but not post-CSD, sociability is rank-dependent. Change of social preference in rank 1 mice was significantly greater after CSD compared to rank 4 mice. (A) Timeline of behavioural studies investigating the effect of dominance status on susceptibility to chronic social stress. The age of mice is indicated in yellow boxes. (B) Pre-CSDS: Following hierarchy formation, the winning and baseline SI ratios exhibit positive correlation, with dominant mice exhibiting higher sociability. (C) Post-CSDS, winning and SI ratios of ranked mice did not differ significantly. (D) Rank 1 displayed a higher change in SI ratio following CSDS. (E) Rank 1 and rank 4 mice do not exhibit anhedonia in the SPT. (F) Daily wheel-running activity profiles are similar across ranks. (G-H) There were no significant differences between groups in either trait or state anxiety tests. N_rank 1_=16, N_rank 4_=15. Data are shown as mean ± SEM. *p<0.05. CSDS: chronic social defeat stress. SI: social interaction. SPT: sucrose preference test TT: tube test. UMA: urine marking assay. ABT: agonistic behaviour test. WST: warm spot test. OF: open field. EPM: elevated plus maze.

#### 3.3.1 Social dominance could be a predictor of sociability but not of stress-susceptibility

Following the two weeks of hierarchy establishment and maintenance, rank 4 (subordinate) mice had a significantly lower SI ratio than the rank 1 (dominant) mice **Fig.3B** left: mean difference = 0.21±0.09; Unpaired t-test t(29)=2.374, p=0.0245). Overall, the TT winning ratio positively correlated with the pre-CSDS SI ratio (**Fig.3B** right: Linear regression F(1,54)=4.811, p=0.0326, R^2^=0.08181), with the dominant ranks exhibiting higher sociability than the subordinate ranks. In contrast, after CSDS, there was no difference between the ranks (**Fig.3C** left: mean difference 0.06±0.11; Unpaired t-test t(29)=0.5417, p=0.5922). Additionally, there was no correlation between SI and winning ratio (**Fig.3C** right: Linear regression F(1,54)=0.5259, p=0.4715, R2=0.009645). While both ranks showed a reduction in the SI ratio following CSDS, the change was significantly greater for dominant mice (**Fig.3D**: Unpaired t-test t(29)=2.058, p=0.0487). This raises the possibility that dominant mice are more severely affected by the experience of chronic stress,or that the social interaction ratio of subordinate animals, being lower already at the start, exhibits a floor effect after CSDS. There were no differences in sucrose preference (**Fig.3E**: Unpaired t-test t(16)=0.7069, p=0.4898) or wheel-running activity between ranks after stress exposure (**Fig.3F**: Two-way RM ANOVA stress group effect F(1,22)=0.008877, p=0.9258).

#### 3.3.2 Anxiety profiles were not rank- or stress-experience-dependent

Thigmotaxis profiles in OF and EPM were not different between ranks 1 and 4 either before or after CSDS (**Fig.3G**: Unpaired t-test OF: t(29)=0.2184, p=0.8287; EPM: t(29)=0.03974, p=0.9686; **Fig.3H**: Unpaired t-test OF: t(29)=0.2516, p=0.8031; EPM t(29)=0.3931, p=0.6971). Measures of explorative and locomotive behaviour were also comparable in all cases (**Fig.S2**). Hence, we cannot report any rank-dependent differences in anxiety measures.

## 4. Discussion

### 4.1 Chronic exposure to social defeat did not render mice subordinate

We anticipated that CSDS exposure may have differential effects on hierarchy formation such that susceptible mice may be more prone to subordination as they exhibit compromised ability to handle stressful conflict situations. In contrast, we predicted that resilient mice may exhibit dominant status as they may have acquired a more adaptive strategy for adjusting to new social cohorts, which enabled them to overcome any putative adverse effects of CSDS. However, our findings do not show any clear effect of prior stress exposure on hierarchy formation. While depression is mainly associated with despondency and social withdrawal, aggression is also a common symptom in human depressive states [39]. Moreover, the chronic unpredictable stress model was reported to increase aggression, hostility, and social dominance in rodents [40]. Our observation that both susceptible and resilient mice exhibit the same amounts of dominance or subordinate status as evidenced by the equal distribution of the winning ratio suggests that exposure to CSDS does not induce aggression or other behavioural adaptations related to hierarchy formation. Our study did not quantify push, retreat, and resistance behaviours of animals within the tube so we cannot exclude the possibility that CSDS-exposed mice win via a more or less effortful strategy than the stress-naïve controls, for example by “freezing” in the tube instead of pushing until the opponent retreats. Additional measures of animals’ agonistic propensity could delineate whether increased aggression would account for these observations. To further determine (i) the effects of stress on susceptible/resilient mice and (ii) measure the differences in dominant and subordinate mice, we investigated whether diurnal activity was disrupted following CSDS exposure. Since a number of studies report that stress affects circadian rhythms and sleep-wake cycle [41] we predicted that stress exposure following CSDS would lead to disruptions in daily rhythms. Daily wheel running activity is a standard measure of internal rhythms where mice will typically exhibit low activity in the day and high activity at night. We did not observe any obvious effect of CSDS on total activity counts in daily wheel running activity since both susceptible and resilient mice exhibit similar rhythms to stress naïve mice. While we report measures of sociability, anxiety, and locomotive behaviour, it is nonetheless difficult to account for the complete array of side-effects that CSDS may have on cognition and physiology, and how these in turn may affect social dominance. Though our initial study measured the effect of stress on winning ratios in completely new social groups, another valuable question would be to examine how stress would affect integration of an individual in an already established group of stress-naïve conspecifics.

### 4.2 Dominant and subordinate mice are equally susceptible to the adverse effects of chronic social stress

The differences in social competitiveness were not related to the overall differences in stress-susceptibility. However, subordinate, but not dominant, mice exhibited decreased average baseline social preference prior to CSDS. Recent observations that subordinate mice exhibit changes in slow-wave sleep (SWS) and rapid eye movement (REM) activity may be indicative of experience to aggression during hierarchy formation [16]. Thus, the decreased social preference prior to CSDS we observe in subordinate mice may be indicative of social stressors experienced by these groups during hierarchy formation. This coincides with recent suggestions of high levels of intrinsic stress in in selectively bred socially-submissive mice [42]. Moreover, our observation of a positive correlation between wining ratio and social interaction ratio further highlights the association between hierarchy status and social preference. However, these differences in social preference between dominant and subordinate groups disappears after CSDS exposure as both hierarchical groups exhibit susceptible phenotypes as evidenced by the similar low social interaction scores. It was recently reported that dominant mice exhibited increased sleep fragmentation, where it was proposed that the stressful effects of constantly maintaining the dominant status lead to fragmentated sleep [16]. Since sleep and circadian rhythms are intimately linked, we wondered whether dominant and subordinate mice would exhibit different daily rhythms following stress exposure. Analysis of daily wheel running activity did not show any difference in diurnal total activity counts per hour in dominant and subordinate mice.

A recent study reported that dominant males are more susceptible to CSDS [26] while we find that dominant mice exhibit the largest change in sociability when we measure SI scores before and after CSDS when compared to subordinates. Our data did not completely replicate the earlier study [26], however, several conditions differ between our two studies. The previous paper used five weeks of cohabitation prior to dominance testing and CSDS stress, while we opted for a two-week period as it was shown to be sufficient for hierarchy formation and is more commonly used in the majority of social dominance studies. Therefore, the inconsistency in stress-susceptibility may arise because the effects of group-housing are relatively mild at two weeks or because more profound effects are only observable after prolonged occupation of a rank. Furthermore, another recent study reported subordinate mice having higher depressive-like behaviour, as well as hormonal and expression levels of genes associated with stress, after only two weeks of group-housing [7]. Therefore, one hypothesis may be that the dominants suffer more severe stress when having to maintain their position throughout a longer period of time, while during the initial establishment of hierarchy subordinates experience more stress. Our findings support this hypothesis since subordinate mice already exhibit a susceptible-like sociability phenotype prior to introduction of any additional stressors as evidenced by the significantly lower pre-CSDS SI ratio. As baseline SI measurements were not reported in the study by Larrieu et al. [26] we cannot make a direct comparison with hierarchies maintained for a longer period.

It is possible that increased vulnerability to chronic stressors may be an effect of long-term dominance, arising from the struggles to maintain the rank position, similar to how long-term subordination in the visible-burrow system induces a stress-phenotype [28]. Nonetheless, studies on hierarchy maintenance in mice showed that there was a large degree of variability between social groups in overall stability, time taken in establishing the hierarchy and in the degree of despotism of the alpha male [43]. As a result, we would expect rank-related differences to arise over a variable timescale, making duration of group-housing a significant contributor to the effects of social hierarchies on behaviour and physiology. Moreover, another explanation for the differences between the studies may be due to the type and strength of stressors used. Acute and chronic stressors can have very different effects on neurophysiological function where for example strong stressors lead to increased firing, while longer term weaker stressors induced decreased firing, in the brain reward circuits [44–46]. Thus, differences in the stress paradigm used in these studies likely induced different changes in neural circuits that will have different effects on behavioural processes such as motivation and aggression during hierarchy formation resulting in differences in observed responses. [36].

## 5. Conclusions

In summary, our results suggest that susceptibility or resilience to prior exposure to chronic social stressors does not determine social status. In contrast, the continuous stress of establishing and maintaining hierarchy has differential effects on mice of distinct ranks. In newly established groups, low-status, subordinate, mice exhibited lower preference to novel social targets, but exposure of the social group to chronic social stress had a greater effect on the sociability of high-status animals.

The clinical consequence of social stress is increasing as the number people living in urban settings increases together with modern life-work demands. Thus, there is a need to expand our knowledge of stress-related factors influencing social behaviours for the purposes of developing appropriate therapeutics. Refinement of animal models that currently assumes that shared housing implies greater phenotypic similarity though recent studies showing social dominance accounted for more variation in mice than cage-identity [47] would be useful to systematically study the behavioural and physiological consequence of social stress.

## Author contributions

**Merima Sabanovic** Conceptualization, Methodology, Investigation, Formal analysis, Visualization, Writing - Original Draft and Review & Editing. **He Liu**: Methodology, Investigation, Supervision. **Vongai Mlambo**: Validation, Investigation, Writing - Review & Editing. **Hala Aqel**: Investigation. **Dipesh Chaudhury**: Conceptualization, Funding Acquisition, Writing - Review & Editing.

## Acknowledgements

The authors acknowledge the kind assistance of William Presley and Fatin Qamash for maintaining the animals and Ayesha Al Hammadi for help with performing chronic social defeat and tube testing, and other members of the Chaudhury lab for advice on experiments.

## Data availability

The authors declare that the data supporting the findings of this study are available on request from the corresponding author (D.C.).

## Supplementary Figure Legend

**Fig. S1.**
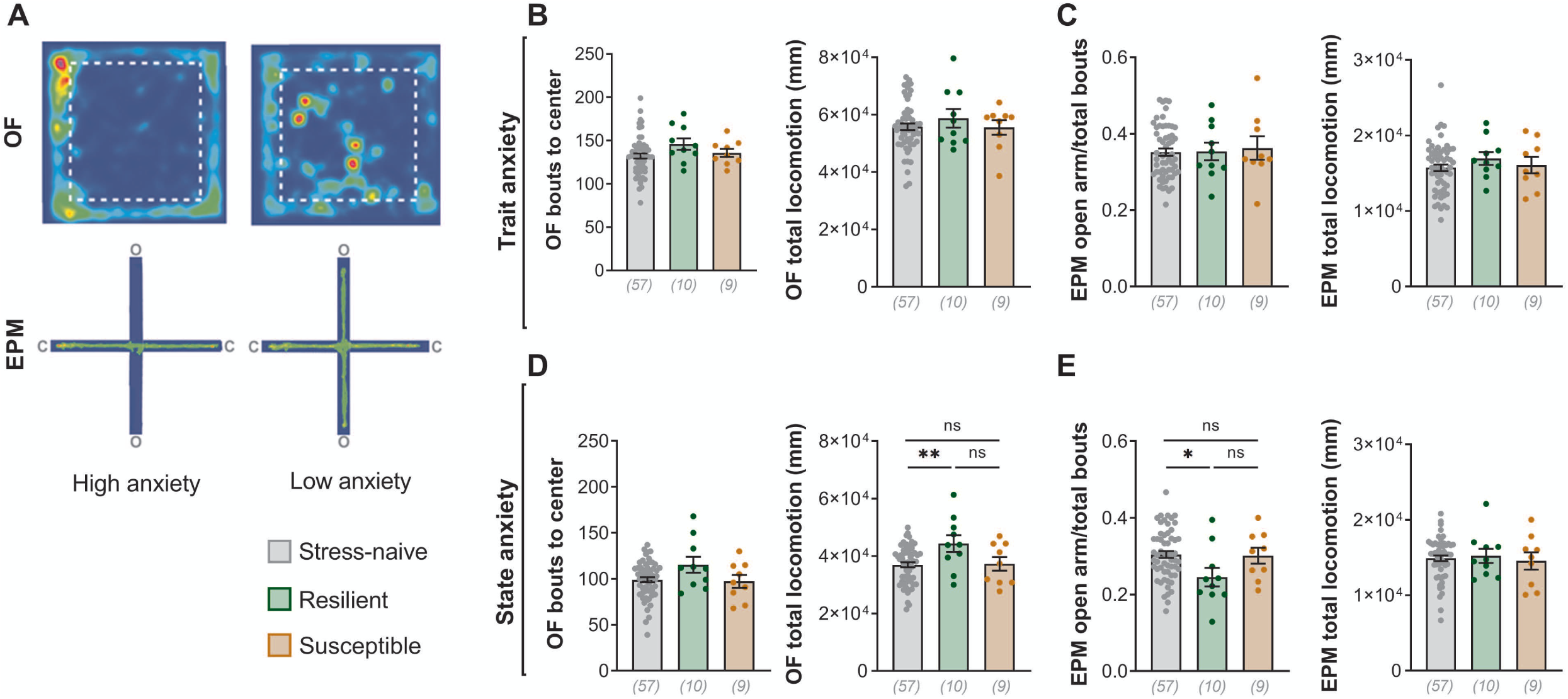
(A) Representative trace from the anxiety tests. Mice with a higher anxiety-like profile remain closer to walls (in OF) or inside closed maze arms (in EPM) for longer compared to the less anxious mouse. O=open arms. C=closed arms. Stress groups do not differ in their exploratory behavioural profiles in trait OF (B) (bouts: One-way ANOVA F(2,73)=1.714, p=0.1873; locomotion: Kruskal-Wallis H(2)=3.853, p=0.8248); or EPM (C) (One-way ANOVA bouts: F(2,73)=0.08528, p=0.9183; locomotion: F(2,73)=0.6106, p=0.5458). (D) In state OF, while the number of bouts is the same across groups, resilient mice cover more distance as compared to the stress-naive but not susceptible mice. One-way ANOVA bouts: F(2,73)=2.671, p=0.0759; locomotion: F(2,73)=4.617, p=0.0129, post-hoc Tukey naive-resilient p=0.0095, naive-susceptible p=0.9910, resilient-susceptible p=0.0858. (E) In state EPM, locomotive behaviour is similar between groups, but resilient mice exhibit reduced number of open arm entries as compared to the stress-naive controls, but not susceptible mice. Bouts One-way ANOVA F(2,73)=3.234, p=0.0451, post-hoc Tukey naive-resilient p=0.0356, naive-susceptible p=0.9921, resilient-susceptible p=0.1783; locomotion Kruskal-Wallis H(2)=0.1805, p=0.9137. Data are shown as mean ± SEM. *p<0.05. N numbers given in brackets.

**Fig. S2.**
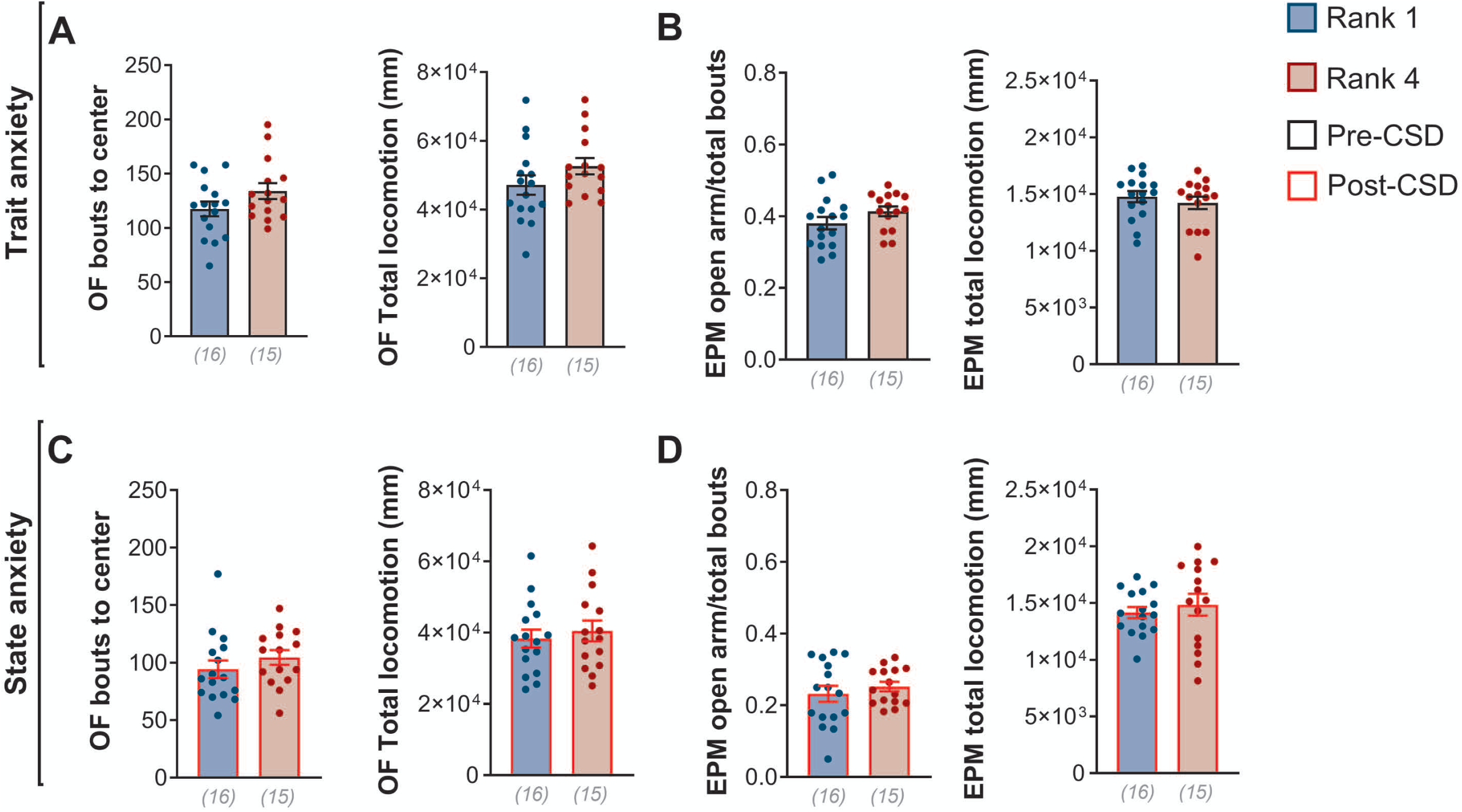
Exploratory profiles of ranks are not different in either trait or state anxiety test. (A) Unpaired t-test bouts: t(29)=1.662, p=0.1072; locomotion: t(29)=1.474, p=0.1514. (B) Unpaired t-test bouts: t(29)=1.493, p=0.1461; locomotion: t(29)=0.7439, p=0.4629. bouts: Mann-Whitney U=81, p=0.1266; locomotion: Unpaired t-test t(29)=0.5584, p=0.5809. (D) Unpaired t-test bouts: t(29)=0.7688, p=0.4483; locomotion: t(29)=0.652, p=0.5195. Data are shown as mean ± SEM. *p<0.05. N numbers given in brackets.

